# Recent climate change strongly impacted the population dynamic of a North American insect pest species

**DOI:** 10.1101/2024.08.08.607030

**Authors:** Yan Boulanger, Adèle Desaint, Véronique Martel, Maryse Marchand, Salomon Massoda Tonye, Rémi Saint-Amant, Jacques Régnière

## Abstract

Climate change is redefining the dynamics of forest ecosystems globally, particularly through its impact on forest pest populations such as the spruce budworm (SBW, *Choristoneura fumiferana* [Clem.]), a major defoliator in North American boreal forests. This study investigates the shifts in the population dynamics of spruce budworm across its range in response to recent climate change. We used a process-based, temperature-dependent ecophysiological model combined with the ERA5 reanalysis to assess changes in SBW phenology, reproduction rate, winter survival and population growth rates from 1950 to 2022 across North America. Our findings demonstrate a pronounced northward expansion of suitable climate conditions for SBW, accompanied by earlier phenological events and increased reproduction rates in northern regions. Conversely, the southern parts of its range are experiencing increased winter mortality due to warmer temperatures. This study highlights the significant impact of elevated temperatures, particularly during critical developmental windows such as spring and summer, which are pivotal for spruce budworm survival and reproduction. Additionally, our results reveal that the observed shifts in pest dynamics are more strongly driven by climate change than by changes in landscape composition and structure. We estimated that suitable growth rates have shifted northward by over 68 km on average, but this shift reached more than 200 km in the easternmost portions of its range. Climate-induced shift in suitable conditions for SBW underscores the need for adaptive forest management strategies that consider the rapid ecological changes and the potential for increased forest vulnerability due to climatic and biotic stressors. This study provides vital insights that can inform adaptive management ensuring the sustainability of forest ecosystems in the face of ongoing climate change.

## Introduction

Climate change is a threat for global forests, notably for the boreal biome (Gauthier et al., 2015). In Canada, where the boreal forest covers 2.7 M km^2^ (Brandt et al., 2013), climate change-induced increase in temperatures since the last 70 years is twice as important as in the rest of the world (Bush & Lemmen, 2019), putting significant pressure on boreal ecosystems and hence on a large swath of North American forests (Price et al., 2013). Warmer temperature and more severe drought conditions have already been shown to affect boreal forest dynamics through increased tree mortality (Liu et al., 2023; Luo & Chen, 2015; Peng et al., 2011), changes in forest productivity (Boisvenue & Running, 2006; Girardin et al., 2016; Kauppi et al., 2014) as well as regime shifts in natural disturbances (Hanes et al., 2019). Further disruption of boreal forest ecosystem dynamics is projected in the upcoming decades with increasing anthropogenic climate forcing (Boulanger & Puigdevall, 2021). Massive impacts on ecosystem goods and services are to be expected, notably on timber production (Brecka et al., 2020), carbon storage (Dymond et al., 2010; Moreau et al., 2022) and biodiversity (Labadie et al., 2024).

Climate change has important impacts on insect pest outbreaks in northern ecosystems (Duke et al., 2009). As ectotherms, insect development is strongly dependent on temperature variations. Although species responses are complex and vary between seasons and bioclimatic regions (Robinet & Roques, 2010), climate-induced alterations in insect ecophysiological processes can affect voltinism, survival and ultimately population dynamics. For instance, warmer winters have been linked to higher overwintering survival in many insect pest species (Bale & Hayward, 2010; Schneider et al., 2021) while they were also shown to advance species phenology (Huang & Li, 2015). Such climate-induced changes in insect pest population dynamics have already led to shifting or extending range distribution (Jepsen et al., 2008; Sambaraju et al., 2019; Soja et al., 2007, Ma et al., 2021) and is thought to have contributed to modifications in outbreak severity, frequency and extent (Jepsen et al., 2008; Pureswaran et al., 2015; Sambaraju et al., 2019). Further changes in climate conditions could also alter the synchrony between plant and insect phenology, thereby altering the vulnerability of host plants (Ekholm et al., 2020; Pureswaran et al., 2018). Climate-induced disruptions could also alter the enemy-herbivore dynamic by disrupting the ability of parasitoids to affect their pest host species (Stireman III et al., 2005). Projected changes in climate conditions suitable for insect pest species will exacerbate alterations in outbreak regimes (Fleming & Volney, 1995; Logan et al., 2003; Williams & Liebhold, 1995), with varying consequences on forest ecosystems and on the forest sector (Marini et al., 2017).

Spruce budworm (SBW, *Choristoneura fumiferana* [Clem.]) is the most important defoliator of coniferous forests in northeastern North America but outbreaks of this species can occur throughout most of North American boreal forests (see Nealis, 2016 for a review). Outbreaks of this species occur periodically every 30-50 years, can last for more than a decade and can synchronously span over several tens of millions of hectares (Boulanger et al., 2012). Important growth reductions and mass mortality in coniferous host species can occur during outbreaks. This in turn greatly affects forest dynamics while impacting the forest sector and regional economy (Chang et al., 2012). A major outbreak occurred between 1967 and 1991 and spread over 55 Mha over eastern Canada and northeastern USA (Pureswaran et al., 2016). The climate envelope of the species is mostly regulated at its northern edge by the length of the growing season while at its southern edge, it is limited by warm conditions and associated energy depletion during the winter diapause (Régnière et al., 2012). A recent study has shown a northward shift in SBW outbreak distribution during the 20th century (Navarro et al., 2018). Increasing temperatures in the upcoming decades are projected to further affect SBW outbreak dynamics through latitudinal and altitudinal changes in suitable climate conditions (Boulanger et al., 2016; Candau & Fleming 2011; Régnière et al., 2012). Further warming could also increase the synchronicity between host budburst and larvae emergence in spring, as well as affecting the relationship with its parasitoids (Seehausen et al. 2017, 2018). increasing the vulnerability of host species at its northern limits (Pureswaran et al., 2015).

Although changes in climate conditions could severely alter insect pest outbreak dynamics, concurrent modifications in landscapes composition and structure could also play a major role in this respect. Changes in landscape spatial patterns, through alteration in host abundance and connectivity, are known to strongly influence forest pest outbreak dynamics (Cooke, 2024; Cooke et al., 2024; Kausrud et al., 2012). For instance, the abundance of old balsam fir or spruce stands could strongly increase SBW outbreak severity within the mixed and boreal forests of eastern North America (Bouchard et al., 2006; Hennigar et al., 2011; MacLean, 1980). Changes in anthropogenic (Kausrud et al., 2012) or natural disturbance rates (Bouget & Duelli, 2004) could alter forest composition and structure which could have a direct impact on the vulnerability of the landscape to the pest species (Robert et al., 2018). High harvesting rates since the last decades in Quebec have dramatically decreased the abundance of old-growth stands (Martin et al., 2021; Waldron et al., 2020) while increasing the abundance of deciduous pioneer species such as trembling aspen (Dupuis et al., 2020; Marchais et al., 2022). Such a multi decades-long harvesting legacy leading to a concomitant decrease in old coniferous forests, including in SBW hosts, could have altered landscape vulnerability and influenced the dynamics of SBW outbreaks, regardless of changes in climate conditions.

Climate change impact research on insects at a continental scale, including on pest species, has been mostly conducted using deterministic correlative approaches such as species distribution models (e.g., Hill et al., 2016). However, these approaches may fail to address the nonlinear nature of insect responses to climate variables. Also, correlative models cannot identify which and how ecophysiological processes are most likely to be affected by changes in climate conditions (Morin & Thuiller, 2009). Mechanistic weather-driven models can be used to assess impacts on insect development, phenology, survival and ultimately on population dynamics (Lehmann et al., 2018). By simulating adequate insect responses to climate variables, process-based ecophysiological models can give further insights about potential changes in outbreak dynamics than by solely relying on correlative approaches (Morin & Thuiller, 2009).

A SBW outbreak that began in the first decade of 2000s has affected more than 10 million ha in northeastern North America as of 2023 (MRNF, 2023). The rather northern location of the outbreak is puzzling as previous major outbreaks in the 20th century and before occurred much farther south (Bouchard & Auger, 2014; Boulanger et al., 2012; Jardon et al., 2003). Up to now, it is still unclear to what extent recent changes in climate conditions or in landscape vulnerability to SBW could explain such a northward shift in outbreak range. The objective of this study was to assess recent changes (1950-2022) in the suitability of climate conditions for SBW over North America using a process-based temperature-dependent ecophysiological model (Régnière et al., 2012). We assessed climate-induced changes in population dynamics by estimating shifts in insect phenology, reproduction rate, winter survival and potential population growth rates over the last 73 years. We used the state-of-the-art ERA5 reanalysis from ECMWF Copernicus to estimate historical changes in climate conditions driving SBW population dynamics at a continental scale. We further assessed changes in landscape vulnerability by specifically looking at alterations in vulnerable host stand abundance between the last two SBW outbreaks in Quebec. We then estimated the importance of changes in forest landscape vulnerability to SBW compared to trends in weather-driven ecophysiological processes to explain differences between the last two major SBW outbreaks in northeast North America.

### Spruce budworm life history

The spruce budworm completes its annual lifecycle by overwintering as a second instar larva (known as L2), resuming activity in spring to feed on new shoots. After emerging from diapause, the larvae go through four additional stages (from L3 to L6) before pupating, and eventually emerging as adults in mid-summer. Females lay up to 200 eggs the following weeks. Once these eggs have hatched, the larvae quickly find a suitable site to overwinter in a silken shelter. Because they do not feed until the next spring, they subsist on energy reserves from the egg. Temperature is the most important weather variable affecting the development and survival of various SBW larval instars, with warmer temperatures accelerating development time, typically following unimodal non-linear response function (Chuine & Régnière, 2017). The rate of development determines whether the insect can reach its cold-hardy diapausing stage before winter, which is particularly significant at the colder edges of its range (Blais, 1958). In warmer regions, temperature impacts the rate at which diapausing larvae deplete their energy reserves and determines their survival over winter (Han & Bauce, 1997, 2000). Both processes profoundly impact individual odds of survival and hence the species’ population dynamics. Temperature thus plays a crucial role in dictating the severity and synchronization of outbreaks (Peltonen et al., 2002), as well as influencing the distribution of the species and its interactions with host plants (Pureswaran et al., 2015).

## Methods

### The model

We used a spruce budworm seasonality model (Regniere, 1983, 1987, 1990; Regniere & You, 1991), updated with development data for the first instar (L1) from Han et al. (2000). This individual-based model uses the technique of non-linear development summation (Logan et al. 2003) and tracks several hundred individual insects, detailing their development, survival, and oviposition from an initial population of overwintering L2 larvae (L2o). Development rates for each life stage (egg, L1, L2o, L2 to L6, pupa, and adult) are derived from laboratory data across various constant temperatures on an artificial diet. These rates are then used to build a simulation model that accumulates development over time under fluctuating natural temperatures. The model uses a 4-h time step to simulate the “aging” process of insects under variable temperature conditions, with individual development rates adjusted based on temperature, and outputs on a daily time step. Different development rate functions are used for each life stage, with sexes differentiated in the sixth instar and pupal stage. Development rate distributions for all stages, except the first instar, pupa, and adult, are generated from independent random values, with extreme values capped to avoid unrealistic individuals. Oviposition rates are modeled based on the number of eggs remaining in a female and air temperature (Regniere, 1983). Overwintering larval mortality data, influenced by early diapause exposure to warm temperatures, were obtained from Han & Bauce (1997). These data informed a logistic regression model to estimate overwintering survival rates based on pre-storage temperature exposure durations. This model forms the basis of the overwintering survival component, where diapausing larvae draw from a fixed amount of energy at a rate dependent on ambient temperature. From this model, we can then derive larval and adult phenology and assess reproduction rates (defined as the number of L2 progeny produced by the initial L2 in the simulation), winter survival and their product that constitutes potential annual population growth rates. Under cold temperature regimes, exposure of unhatched eggs to frost (temperature < 0°C) is the major additional source of variation in mortality. The model was previously calibrated for the whole SBW distribution and is fully described in Régnière et al. (2012). The model was coded to run in BioSIM 11 (Régnière et al., 2014).

### Data

#### Weather data

We used daily temperatures from the ERA5 reanalysis dataset to assess the annual SBW seasonality, reproduction rates, winter survival and population growth rates. The ERA5 reanalysis dataset is a comprehensive and high-resolution global climate dataset produced by the European Centre for Medium-Range Weather Forecasts (Hersbach et al., 2017). It provides detailed information on various atmospheric, oceanic, and land surface variables from 1940 to the present. ERA5 uses a combination of historical observational data and advanced modeling techniques to generate consistent and accurate records of weather and climate, offering hourly data at a spatial resolution of approximately 31 kilometers (0.25° x 0.25°). We restrained the weather dataset temporally to the 1950-2022 period, and spatially to ERA5 cells covering the entire United States and Canada (Figure 1). Daily ERA5 temperature data were integrated in BioSIM to run the ecophysiological model.

**Fig. 1.**
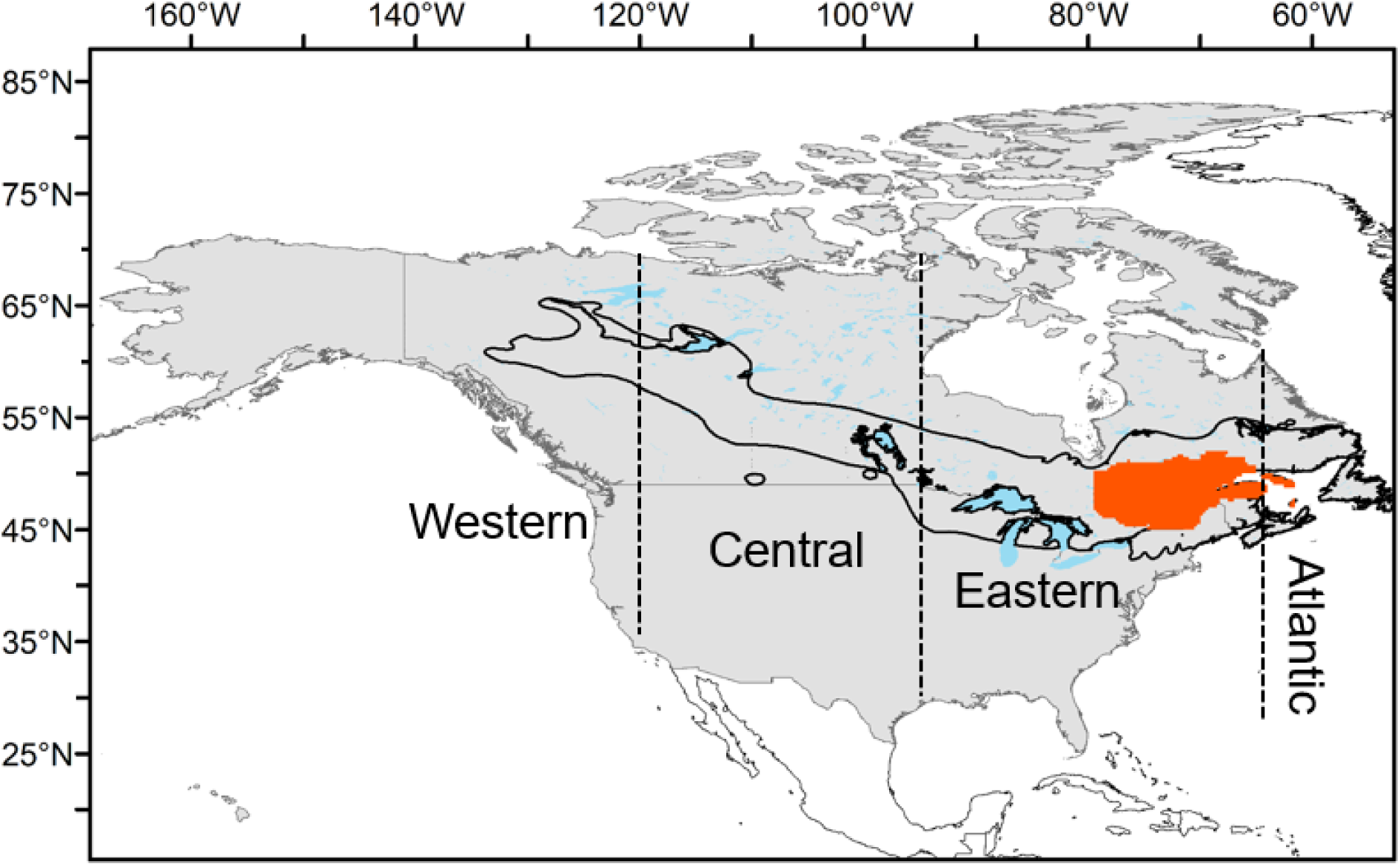
Location of the study area. Cells in gray were analyzed for trends in phenological and ecophysiological parameters for SBW during the 1950-2022 period. Cells in orange were also assessed for changes in SBW landscape vulnerability. The thick black line represents the current SBW range. The delineation of the four regions (Western, Central, Eastern and Atlantic) is also shown by the dashed lines.

#### Forest landscape data

To assess changes in landscape vulnerability to SBW, we focused on data coming from the commercial forests of Quebec. The Quebec province has performed comprehensive and extensive decadal forest inventories within its commercial forest since the mid-1960s (*Ministère des Ressources Naturelles et des Forêts,* [MRNF], 2023). Combined with extensive SBW outbreak severity surveys, this dataset offers a unique opportunity to conduct analyses on recent (< 70 years) changes in forest landscapes and hence in forest landscape vulnerability to SBW between the last two SBW outbreaks. The forest inventory database known as SIFORT (Forest Inventory of the Territory of Quebec) was retrieved from the Open Data portal of the Quebec government (Gouvernement du Québec, 2024; <u>donneesquebec.ca/recherche/dataset/</u>). The SIFORT database compiles forest data in Quebec through a combination of remote sensing technologies, aerial photographs, field surveys, and forest management records. Information is collected at a 14-hectare resolution in ‘tessels’ within which forest composition, age, and other forest data are gathered from the stand located at its centroid. Since 1970, five decadal inventories have been completed. Forest data were compiled differently between inventories, notably regarding stand designation and age class. For this reason, we had to generalize stand composition and age class to the least common denominator between inventories. Stands that were either dominated by host species such as balsam fir and spruces and that were at least 70 years old or considered as “old” in first inventories were considered as highly vulnerable to SBW. The proportion of highly vulnerable stands assessed from the tessels was then reported at the ERA5 cell level.

#### SBW outbreak data

Since 1967, the MRNF has been conducting annual aerial surveys to assess the defoliation caused by the spruce budworm (MRNF, 2023). The aerial surveys are conducted in designated areas, selected on the basis of the extent of defoliation observed in the previous year and predictive surveys of spruce budworm populations. Annual defoliation caused by SBW is categorized into three classes: 1) light, when loss of foliage occurs in the upper third of the crown on a few trees; 2) moderate, when loss of foliage occurs in the upper half of the crown on most trees; and 3) severe, when loss of foliage occurs along the entire length of the crown on most trees. Only foliage loss in crowns of species susceptible to SBW is considered. We only retained years with severe defoliation since lightly defoliated areas could be incomplete in the dataset. If severe defoliation was recorded within the ERA5 cell, the cell was characterized as severely defoliated, indicating that severe defoliation was probable in this area. We then focused on annual defoliation values for the two last SBW outbreaks, i.e., 1967-1991 for the previous major outbreak and 2007-2022 for the ongoing outbreak (Figure 2). For each ERA5 cell, we calculated the number of years in which defoliation was severe during each respective outbreak.

**Figure 2.**
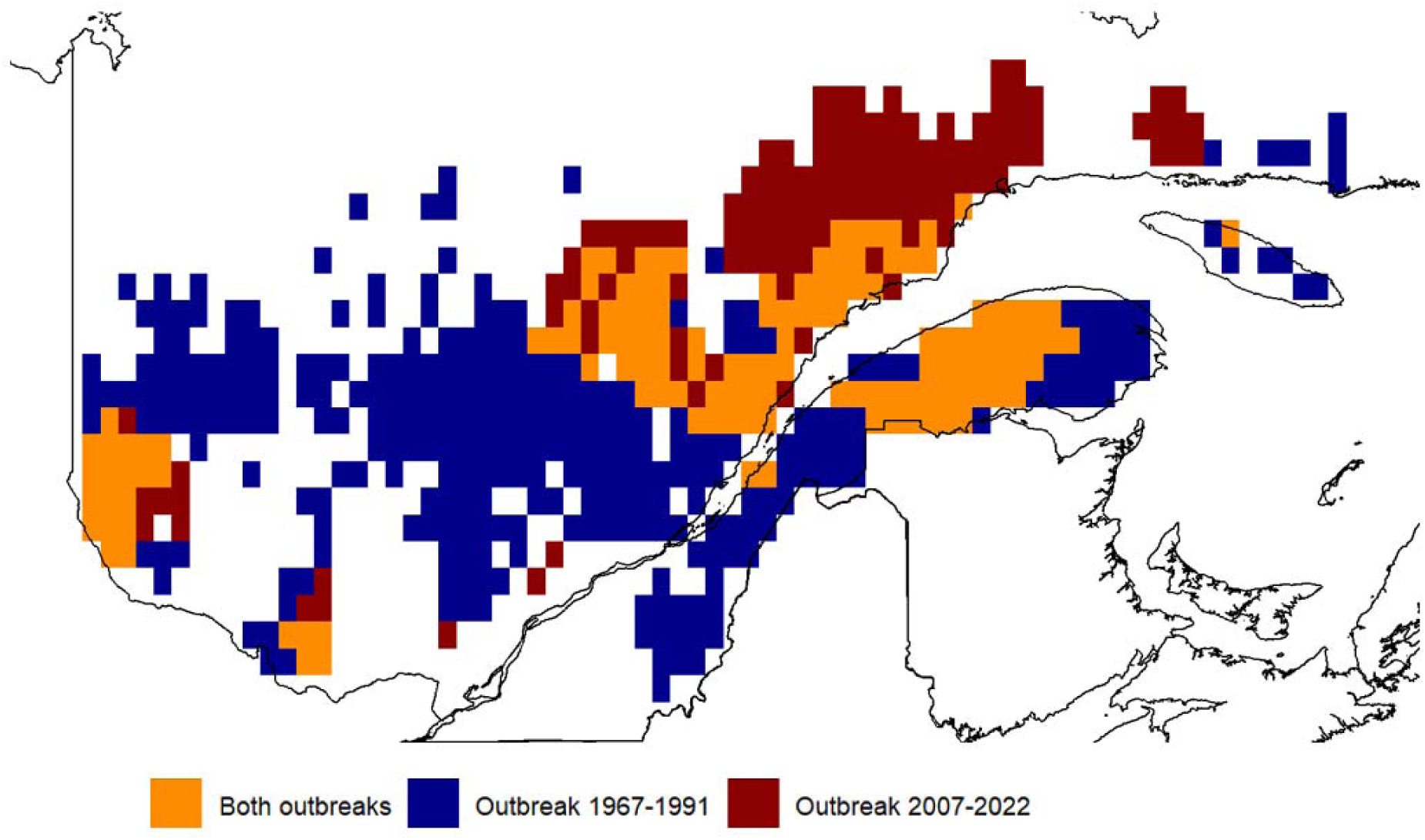
Maps showing the areas affected by at least four years of severe defoliation only during the 1967-1991 (blue), only during the 2007-2022 outbreak (red) or during both outbreaks (orange). Data was summarized at the ERA5 cell resolution across commercial forests in Quebec.

### Analyses

#### Assessing trends in weather, phenology and ecophysiological parameters

For descriptive purposes, we calculated mean temperatures in spring (defined as March, April and May), summer (June, July and August) and fall (September, October and November) for each ERA5 cell throughout the 1950-2022 period. Using daily temperature data for years t and t-1, we then simulated the development of 400 individuals in BioSIM. For each ERA5 cell, we calculated the ordinal date of peak emergence of overwintered L2, L4 and adults for each year of the 1951-2022 period. Using simulation outputs, we assessed the annual reproduction rate and winter survival rate which were both used to calculate the annual population growth rates following Régnière et al. (2012).

We then assessed trends in seasonal temperatures as well as in the phenology and ecophysiological parameters by calculating the Thiel-Sen slope on the 1950-2022 time series at the ERA5 cell level. The Thiel-Sen slope estimator is preferred for assessing temporal trends in climate data because it is robust to outliers as well as being non-parametric, providing reliable and consistent trend estimates without assuming a specific data distribution (Huth & Pokorna, 2004). For all trend analyses, time series were pre-whitened to remove any temporal autocorrelation before assessing slope coefficient and significance. Slopes were considered significant at P IZ 0.1. Thiel-Sen slopes and trends significance were assessed using the *tfpwmk* function in the *modifiedmk* package v1.6 (Patakamuri & O’Brien, 2021) in R v4.3.2 (R Core Team, 2022).

Significant Thiel-Sen slopes for each weather, phenology and ecophysiological parameter were mapped over the study area to delineate potential spatial patterns. As recent climate change was shown to be highly variable along a west-to-east gradient (Bush & Lemmen, 2019), we also further analyzed these trends along a longitudinal gradient by averaging Thiel-Sen slope values for cells at the same longitude. Nonsignificant slopes were assigned a 0 value. To avoid including trends assessed for cells where SBW is not expected to be occurring, we limited this calculation to cells within the current SBW ranges as well as within a 500-km buffer around this range. For graphical and summarizing purposes, we arbitrarily grouped longitudes within four regions, i.e., Western (west of 120°W), Central (120°W - 95°W), Eastern (95°W – 64.5°W) and Atlantic (east of 64.5°W) regions. Trends were reported on a decadal scale as well as over the 1951-2022 period.

We were further interested in assessing the spatial shift in highly suitable conditions for reproduction, winter survival and population growth. Due to the absence of specific thresholds from the Régnière et al. (2012) model for categorizing these ecophysiological parameters as “highly suitable,” we defined such conditions for each parameter as those showing a predicted annual value above the global median using values for the whole period, within the SBW range and its 500-km buffer. Using the Thiel-Sen estimators for each ERA5 cell, we calculated the predicted values for each ecophysiological parameter at both ends of the time series, i.e., in 1951 and in 2022. The total area covered by highly suitable conditions was assessed and compared between both years. Furthermore, for each longitude, we calculated the mean latitude where predicted high suitability conditions were occurring in 1951 compared to 2022. This allowed USA to calculate the predicted latitudinal displacement of highly suitable conditions for each ecophysiological parameter during the 72-yr period.

#### Assessing changes in landscape vulnerability to SBW

To track changes in landscape vulnerability to SBW between the last two major SBW outbreaks, we estimated changes in forest composition and structure as the difference in the proportion of highly vulnerable stands between the first (1970-1983, median year 1976) and fourth (2001-2018, median year 2009) inventories. By doing so, we optimized the timing of the inventories to occur before most of the landscape could be significantly modified by each SBW outbreak per se (see below). Only tessels surveyed during both inventories were analyzed, covering 623,208 km^2^ of ERA5 cells.

#### Assessing the relative importance of climate change and changes in landscape vulnerability

We assessed the relative importance of recent climate change and alterations in forest landscapes on SBW dynamic by comparing trends in ecophysiological parameters as well as in landscape vulnerability in areas heavily defoliated between the last two SBW outbreaks. We first identified ERA5 cells that recorded at least four years of severe defoliation during either outbreak. By doing so, we focused on heavily defoliated areas that were most susceptible to show extensive host mortality (MacLean, 1980). We then identified cells that were heavily defoliated during the 1967-1991 outbreak only, during the 2007-2022 outbreak only and during both outbreaks. For each of these three groups of cells, we reported the 1951-2022 trends in reproduction rates, winter survival and in population growth rates. We further reported changes in landscape vulnerability between the first and fourth decadal inventories for the same three groups. Important differences in trends in ecophysiological parameters or in landscape vulnerability among the three regions could help explain which factors were most important to explain changes in outbreak spatial patterns between the two SBW outbreaks.

## Results

### Trends in seasonal temperatures

Temperatures significantly increased within the SBW range between 1950 and 2022 with the greatest increase occurring during summer (+0.21C per decade), followed by spring (+0.17C per decade) and fall (+0.12C per decade) (Figure 3). Significant increases in summer temperatures occurred all over the SBW range without large spatial discrepancies. Spring temperature increases were most important (∼ +0.2 to +0.3C per decade) in the western and central portions of the SBW range. Spring temperatures did not change significantly over the 1950-2022 period within most of the eastern portion of the SBW range. Most important increases in fall temperature (∼+0.25C to +0.32C per decade) were experienced in the Western region, and at the transition between the Eastern and Atlantic regions, as most of the Central zone did not experience significant changes in fall temperatures (Figure 3).

**Figure 3.**
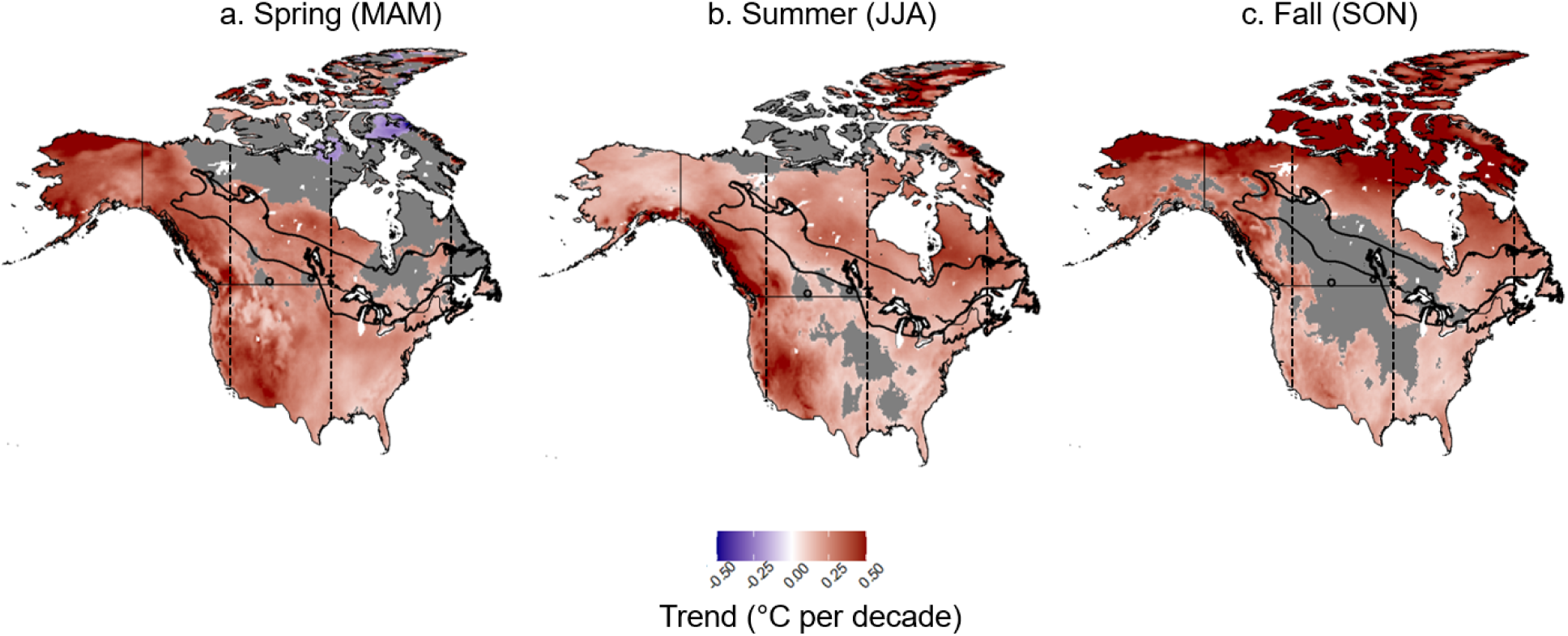
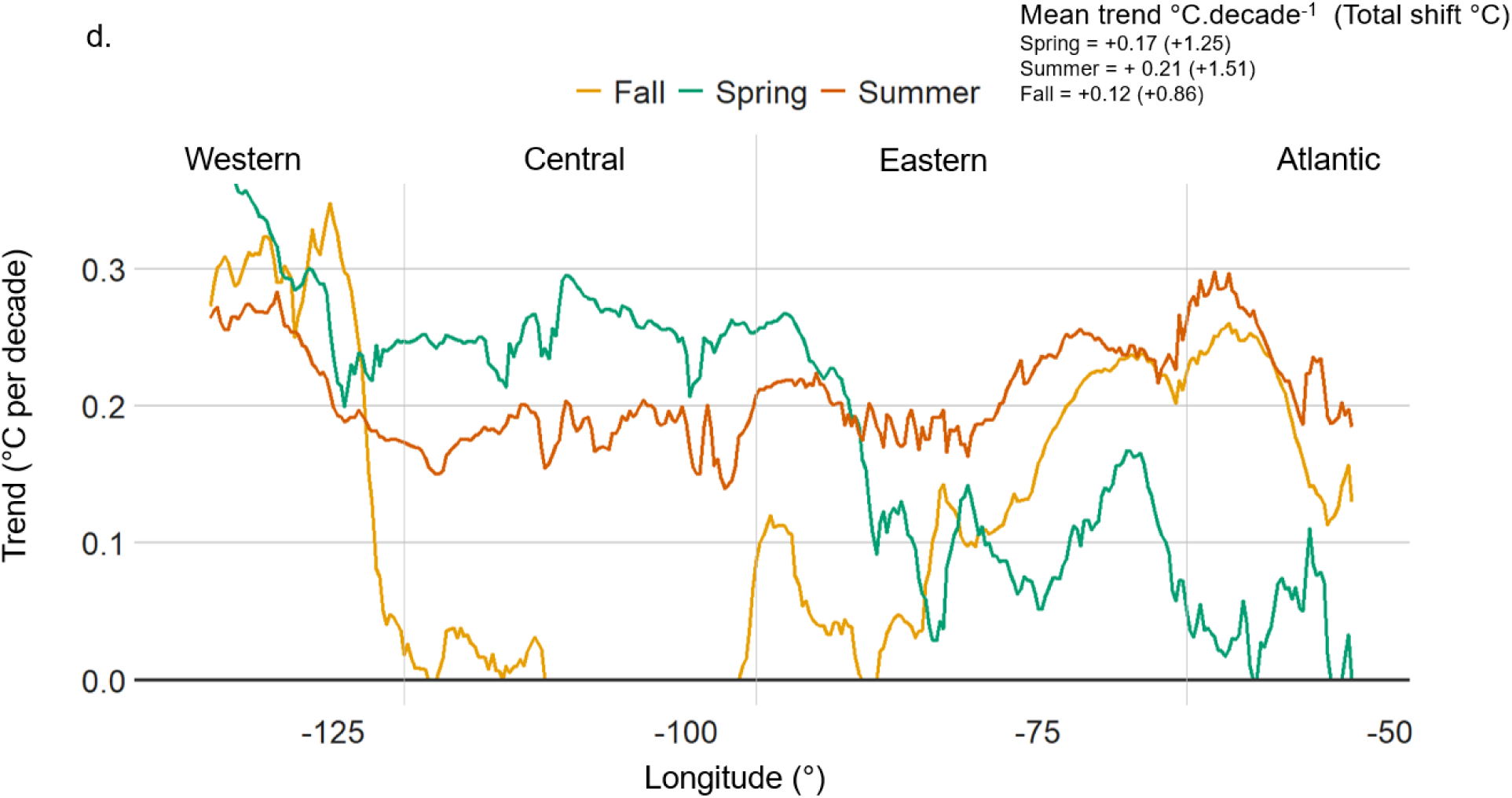
Maps showing decadal trends (°C) in spring (a, MAM: March, April, May), summer (b, JJA: June, July, August) and (c) fall (SON: September, October, November) mean daily temperatures between 1950 and 2022 at the ERA5 cell resolution across North America ERA5 cells showing non-significant trends (P value > 0.1) appear in gray. Western, Central, Eastern and Atlantic regions are arbitrarily delineated at 120°W, 95°W and 64.5°W respectively and are shown as dash lines on maps. (d) Decadal trends for spring (green), summer (dark orange) and fall (light orange) temperatures as averaged by longitude within the spruce budworm range (bold black contour line on a, b and c). Mean decadal trends (and total shift between 1950 and 2022) across the whole spruce budworm range is also shown.

### Trends in SBW phenology

The vast majority of the SBW range experienced significantly earlier larval and adult phenology over the studied period (Figure 4), with trends showing increasingly earlier timings from L2 emergence in spring (−1.11 days per decade) to adult emergence (−1.36 days per decade) on average within the SBW range. Spatial variations in phenology shifts are mirroring changes in spring temperatures between 1950 and 2022. Peak abundances of L2 and L4 occurred up to 20 days earlier in 2022 compared with 1951 in the Western region (Figure 4). In other regions, the timings of L2 and L4 but also adult peak abundances were ∼0.5 to 1.5 days earlier per decade on average along the longitudinal gradient. The timing in adult peak abundance clearly shows a south to north gradient as it occurred *ca* 15 to 30 days (*ca* 2 to 4 days earlier per decade) earlier in 2022 at the northern fringe of the SBW range when compared with predicted values in 1951. In Central regions, most important trends in the timing of adult peak abundance occurred north of the current SBW range.

**Figure 4.**
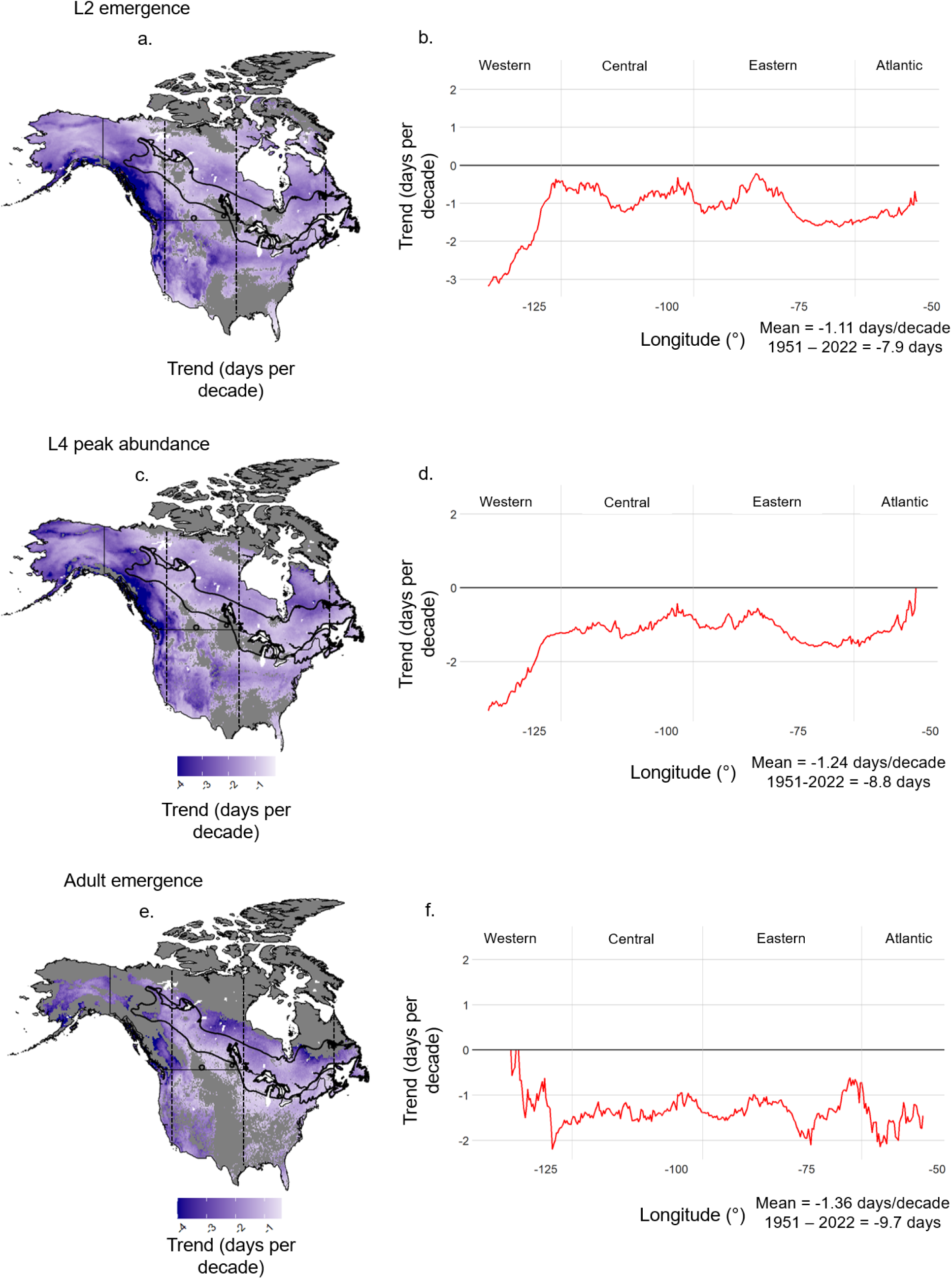
Maps showing decadal phenology trends (in days) in peak L2 emergence (a), peak L4 abundance (c) and peak adult emergence (e) between 1951 and 2022 at the ERA5 cell resolution across North America ERA5 cells showing non-significant trends (P value > 0.1) appear in gray. Western, Central, Eastern and Atlantic regions are arbitrarily delineated at 120°W, 95°W and 64.5°W respectively and are shown as dash lines on maps. Decadal trends for these parameters as averaged by longitude within the spruce budworm range (bold black contour line on a, c and e) is also shown in b, d and f respectively. Mean decadal trends (and total phenology shift between 1951 and 2022) across the whole spruce budworm range is also shown. Negative values mean earlier phenology.

### Trends in reproduction rate, winter survival and in population growth rates

Within most of the SBW range, reproduction rates did not change significantly, except in northern portions of the Eastern and Atlantic regions, as well as in the westernmost tip of the SBW range (Figure 5). As opposed, winter survival significantly decreased virtually everywhere within the SBW range except for a very small portion of the Eastern and Atlantic regions. When combining reproduction rates and winter survival, simulated population growth rates generally declined over much of the SBW range. However, mirroring trends in reproduction rates, population growth rates increased considerably in the northern tier of the Eastern and Atlantic regions.

**Figure 5.**
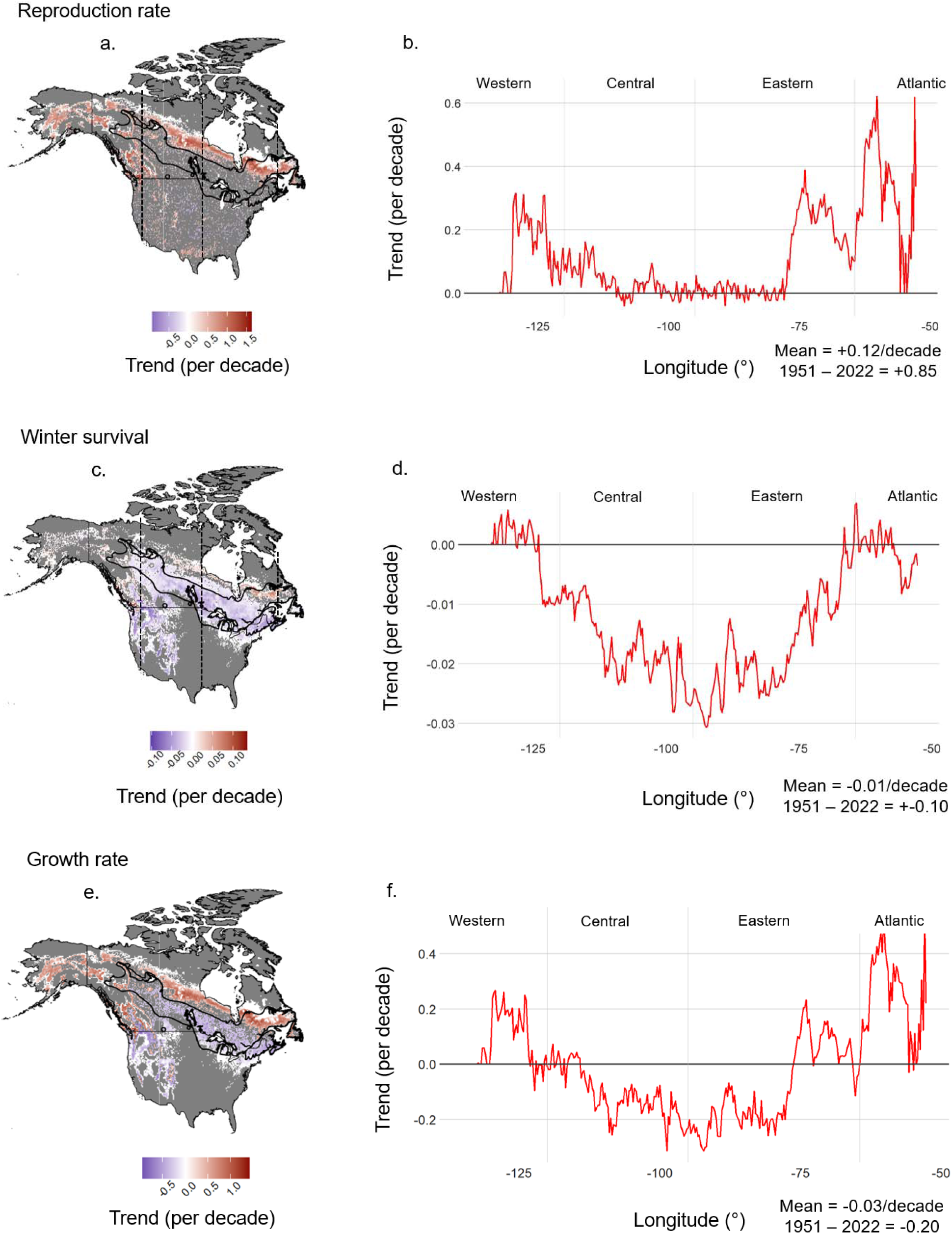
Maps showing decadal trends in reproduction rate (a), winter survival (c) and population growth rate (e) between 1951 and 2022 at the ERA5 cell resolution across North America ERA5 cells showing non-significant trends (P value > 0.1) appear in gray. Western, Central, Eastern and Atlantic regions are arbitrarily delineated at 120°W, 95°W and 64.5°W respectively and are shown as dash lines on maps. Decadal trends for these parameters as averaged by longitude within the spruce budworm range (bold black contour line on a, c and e) is also shown in b, d and f respectively. Mean decadal trends (and total shift between 1951 and 2022) across the whole spruce budworm range are also shown.

As a result, highly suitable areas for reproduction rates shifted northward with new areas in 2022 appearing along a 30-100 km band north of conditions that were highly suitable for reproduction in 1951 (Figure 6). These new areas appeared mostly outside the current SBW range except in the Eastern and Atlantic regions. In these regions, highly suitable conditions for reproduction shifted 100-300 km north. This shift is much larger than the average of 38 km within the continental SBW range. Regions with highly suitable winter survival shifted more strongly to the north, averaging 78 km at the continental scale mostly because of strong contraction of these conditions at the southern trailing edge. Such a contraction resulted in a 200-400 km northward shift in conditions highly suitable for winter survival in the Eastern and Atlantic region (Figure 6). Regions with highly suitable population growth rates shifted an average 68 km north continentally. Largest shifts within the SBW range occurred in the Eastern and Atlantic regions where they shifted 200 to almost 400 km north. As for reproduction rates, new regions with highly suitable growth rates appeared north of the current SBW range in the Central and Western regions. When considering these spatial shifts, the total area highly suitable for reproduction rates increased by 17% between 1951 and 2022 while areas highly suitable for winter survival decreased by 34% (Figure 6). Overall, the total areas showing highly suitable population growth rates remained virtually the same between 1951 and 2022 (+1%).

**Figure 6.**
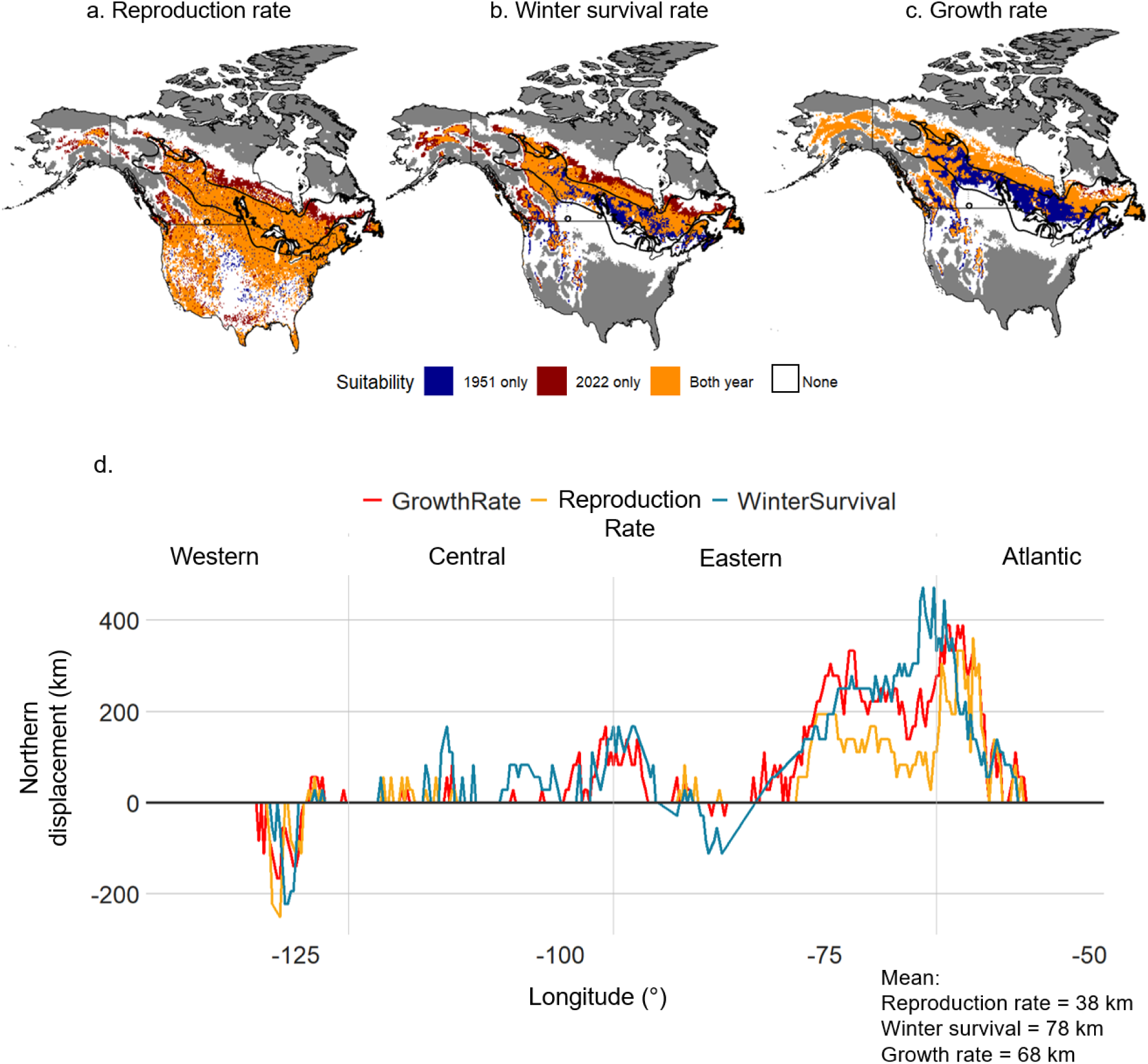
Maps showing the displacement of highly suitable areas for (a), reproduction rate (b) winter survival and (c) population growth rate between 1951 and 2022 at the ERA5 cell resolution across North America. Highly suitable area for each parameter were defined as those showing a predicted annual value above the global median using values for the whole period, within the SBW range and its 500-km buffer. Areas that were suitable only in 1951 are shown in dark blue, areas that were highly suitable in both 1951 and 2022 are shown in orange, areas that were suitable only in 2022 are shown in dark red and areas that were not highly suitable during both years are shown in white. Northern displacement for highly suitable areas as averaged by longitude within the spruce budworm range (bold black contour line on a, c and e) is also shown in d. Western, Central, Eastern and Atlantic regions are arbitrarily delineated at 120°W, 95°W and 64.5°W respectively. Total northward displacement between 1951 and 2022 for winter survival, reproduction rate and population growth rate across the whole spruce budworm range is also shown.

### Importance of recent climate change and changes in landscape vulnerability on SBW outbreaks

Severe defoliation associated with the ongoing outbreak is located on average *ca* 110 km (49.1 ± 1.2°N, mean ± SD) north of areas that were severely defoliated during the 1967-1991 outbreak (48.1 ± 1.0°N, mean ±SD) (Figure 2). There is a clear south to north trend in areas severely affected by SBW between the two outbreaks. Areas severely affected only in the 2007-2022 period are on average 200 km north (49.8 ± 1.2°N, mean ±SD) than those severely affected only by the 1967-1991 outbreak (48.0 ± 1.0°N, mean ±SD). Areas affected by both outbreaks are located midway on average at 48.5° ± 0.9°N (mean ±SD).

Abundance of old balsam fir/spruce stands in Quebec commercial forests have generally declined (−4.2 ± 17%) between the first and fourth inventories (Figure 7). Decreases are generally similar between areas severely affected by SBW by either outbreak. Yet, the variance in changes in proportion of old balsam fir/spruce stands increases from south to north with more important ERA5 cell-level decreases and increases generally observed within areas that were severely affected only by the ongoing outbreak (Figure 7).

**Figure 7.**
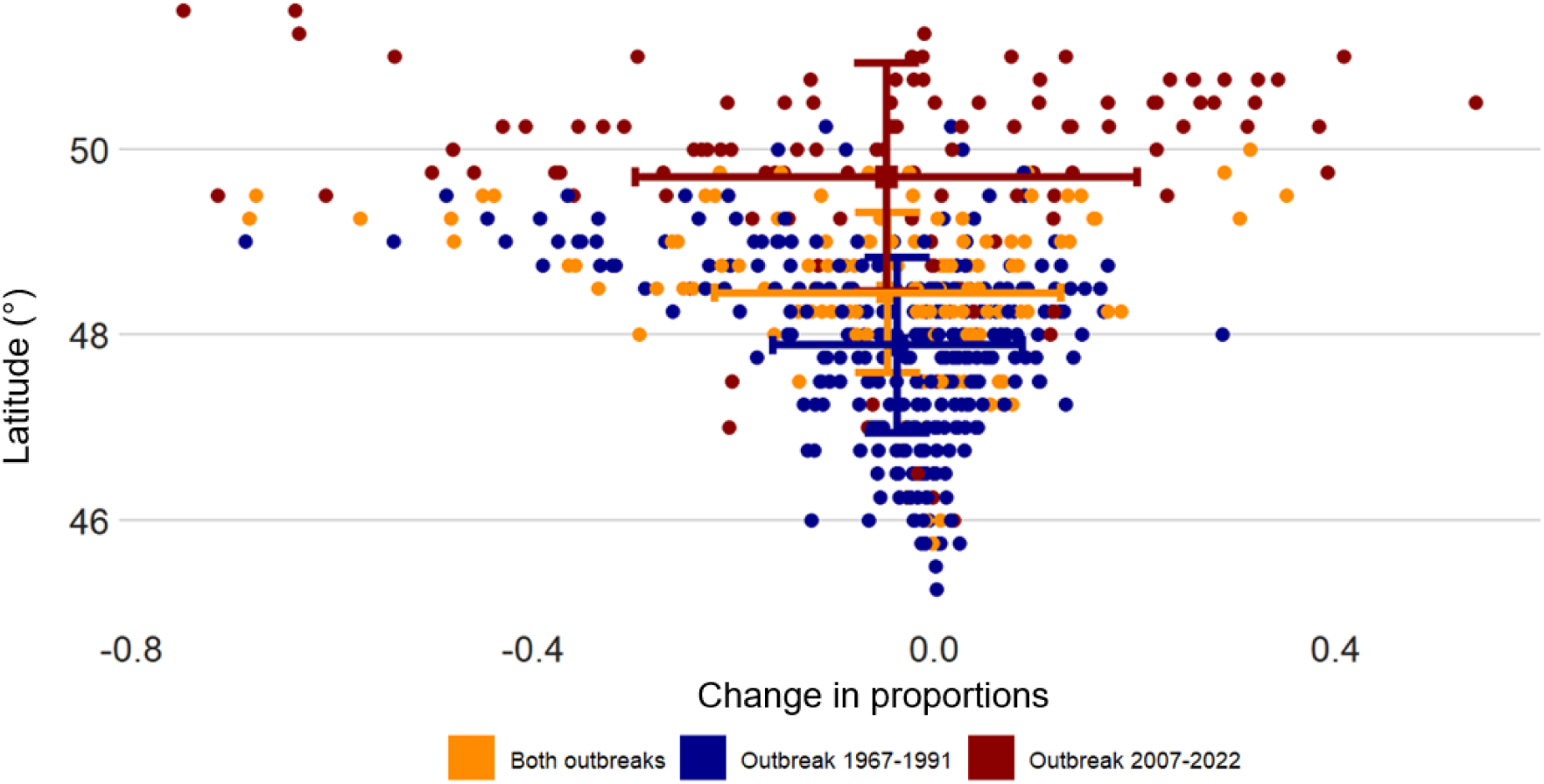
Changes in proportion of old balsam fir and spruce stands between the first (median year: 1976) and fourth (median year: 2009) forest inventories at the ERA5 cell resolution across commercial forests of Quebec according to latitude within areas affected by at least four years of severe defoliation only during the 1967-1991 (blue), only during the 2007-2022 outbreak (red) or during both outbreaks (orange). The centroid for each group of affected regions is also shown as well as its x and y standard deviation.

Differences in trends in ecophysiological processes are more pronounced between areas severely affected by the two outbreaks (Figure 8). Trends in reproduction rates, winter survival and population growth rates were on average much higher in areas severely defoliated only in 2007-2022. On average, trends in reproduction rates were barely positive in areas severely defoliated only in 1967-1991 (0.07 ± 0.21) and during both outbreaks (0.10 ± 0.23) whereas reproduction rates have strongly increased (0.50 ± 0.38) in areas that were severely defoliated in the ongoing outbreak (Figure 8). Likewise, winter survival has barely declined in regions severely defoliated only between 2007 and 2022 whereas ERA5 cells showing decline in winter survival are much more important in areas affected by both outbreaks or only between 1967 and 1991. As a result, ERA5 cells showing positive population growth rates are much more common in areas severely affected by the ongoing outbreak (66%) than in areas only severely affected by the outbreak in the 20th century (10%).

**Figure 8.**
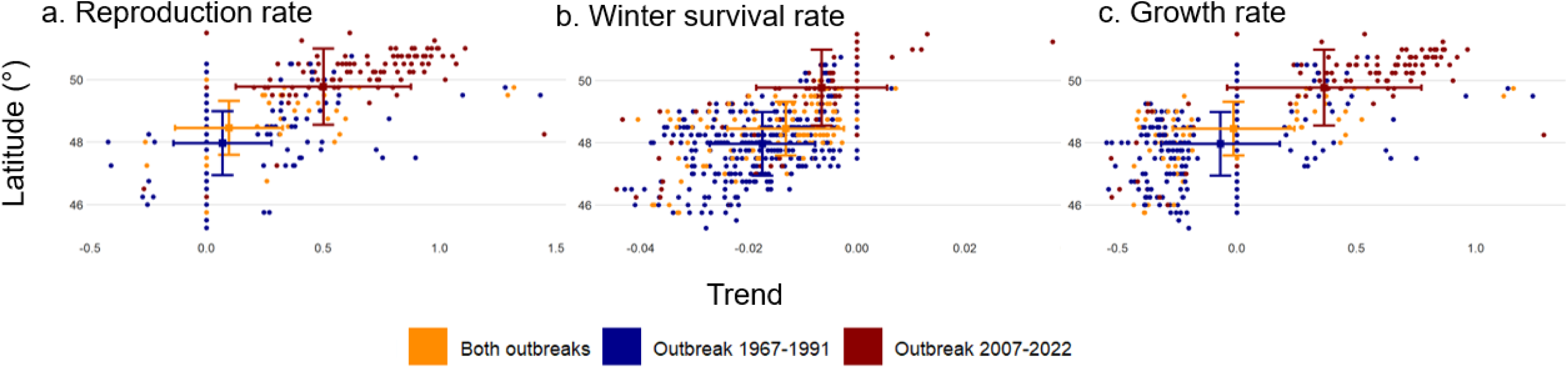
Decadal trends in (a), reproduction rate (b) winter survival and (c) population growth rate between 1951 and 2022 at the ERA5 cell resolution across commercial forests of Quebec according to latitude within areas affected by at least four years of severe defoliation only during the 1967-1991 (blue), only during the 2007-2022 outbreak (red) or during both outbreaks (orange). The centroid for each group of affected regions is also shown as well as its x and y standard deviation.

## Discussion

Our results show that recent climate change strongly altered the phenology of the SBW and shifted the climate envelope of suitable conditions for SBW considerably northward. Rather than using a correlational approach, the use of a mechanistic weather-driven model allowed USA to better identify which ecophysiological processes have been altered by recent climate change, thereby changing the insect pest climate envelope. Overall modifications in winter survival and reproduction rates resulted in an average northward shift of 68 km in regions with highly suitable population growth rates, with shifts exceeding 200 km in eastern and Atlantic North America Northward shifts in climate conditions suitable for SBW align with those recorded from other studies performed on forest-associated insect pest species in the northern hemisphere. For example, the expansion of the southern pine beetle, *Dendroctonus frontalis* Zimmermann (Coleoptera: Curculionidae) in northern states in the USA in recent decades represents a significant northward shift of its historical range of 85 km per decade since 2002 (Lesk et al., 2017). Likewise, the northern limit of the range of the mountain pine beetle, *Dendroctonus ponderosae* (Hopkins) (Coleoptera: Curculionidae), has shifted northward by approximately 391 km over a 26-year period (1965-1996) in western Canada (Sambaraju et al., 2019). In Europe, the Spongy moth’s, *Lymantria dispar dispar* (L.) (Lepidoptera: Erebidae), range has expanded to the 61st parallel, with an average spread rate of 50 km/year while the Nun moth, *Lymantria monacha* (L.) (Lepidoptera: Erebidae), has expanded its range northward by approximately 200 km in Finland over the past two decades (Fält-Nardmann et al., 2018). Warmer winter temperatures (Lesk et al., 2017), increased survival rates in cold climate (Fält-Nardmann et al., 2018), phenological synchronization with host plants (Jepsen et al., 2008, 2011), extended growing season (Sambaraju et al., 2019), and developmental plasticity (Ponomarev et al., 2023) were all cited as factors driving the climate-driven northward expansion of insect pest species.

Recent climate changes are causing phenology to occur much earlier across the entire SBW range. Similarly, climate change triggered earlier phenology, resulting in earlier swarming periods and more generations per year for the European Spruce Bark Beetle, *Ips typographus* (L.) (Coleoptera: Curculionidae), (Berec et al., 2013; Jacoby et al., 2019). Likewise, warmer conditions shifted both Nun moth and Spongy moth phenology earlier in northern Europe (Fält-Nardmann et al., 2018) while earlier development stages due to accumulated temperatures probably enhanced outbreak severity and the northward expansion of the Hemlock Woolly Adelgid, *Adelges tsugae* Annand (Hemiptera: Adelgidae), in northeastern USA (Crimmins et al., 2020). We show that earlier phenology is allowing the SBW to more easily complete its life cycle before winter in the northern portions of its range, boosting reproduction rates and expanding the area suitable for population growth northward. Although not assessed in the current study, earlier phenology could also contribute to increased synchrony with previously less vulnerable hosts, such as black spruce, *Picea mariana* (Mill.) (Pinaceae), which would contribute to further increase population growth rates northward (Bellamin-Noël et al., 2021; Pureswaran et al., 2018). Increasingly warmer temperatures during diapause in the southern part of SBW range is contributing to contracting the southern trailing edge of highly suitable conditions southward. Warm temperatures during the diapause of L2 larvae are linked to important energy depletion during a period when the larvae can not feed, leading to higher mortality (Han & Bauce, 1997). Deleterious impacts of warm winters on SBW contrasts with other insect species whose recent or future range expansion have been linked to climate change-induced warmer winters in recent decades (Duan et al. 2020; Régnière & Bentz, 2007; Lesk et al., 2017; McAvoy et al., 2017; Samabaraju et al., 2012). The SBW is known to be very cold-tolerant (Delisle et al., 2022) and as such warmer extreme minimum temperatures barely affect the northern expansion of its suitable climate conditions. Such “decoupling” between the pest species and its thermal environment represents a phenomenon that is projected to become more common in the next decades with increased anthropogenic climate change (Bale & Hayward, 2010).

The general northward shifts in highly suitable conditions for population growth could have more important impacts on forest succession in the Eastern and Atlantic regions than in Western and Central regions. Much higher reproduction rates, and hence population growth rates, in northern portions of the Eastern and Atlantic regions occurred where its most vulnerable host, balsam fir, is a dominant or codominant species (Beaudoin et al., 2014; De Grandpré et al., 2018). However, newly available and highly suitable areas for population growth in Central and Western regions emerged outside the current SBW range, in areas where most vulnerable host species are sparsely distributed (Beaudoin et al., 2014). Simultaneously, these regions are already, and increasingly, exposed to wildfires under increasing anthropogenic climate forcing (Hanes et al., 2019). This natural disturbance helps make forest landscapes less vulnerable to SBW by promoting the growth of young and less favorable pioneer tree species, such as jack pine and trembling aspen. Recent and future shifts in the SBW climate envelope should thus increase the mismatch with its host in Western and Central Canada, further decreasing the potential for severe outbreaks in this area (Boulanger et al., 2016).

Recent shifts in climate conditions favorable for SBW population growth, rather than changes in forest landscapes, are likely to have affected the recent SBW outbreak dynamic in northeastern North America Since the first decade of the 2000s, a growing SBW outbreak has affected more than 10 Mha in the commercial forests of Quebec. This outbreak spread from foci located mostly within the northern portions of the mixed forests as well as within the boreal forests (Bouchard & Auger, 2014). Although the current outbreak is not over as of 2023, very sparse severe defoliation associated with the ongoing outbreak has occurred south of 47°N (MRNF, 2023). This is a rather unusual situation when compared to previous outbreaks over at least the last 500 years in eastern Canada (Boulanger et al., 2012; Jardon et al., 2003). We found that the severe defoliation associated with the ongoing outbreak has occurred, on average, 110 km farther north than during the previous outbreak. The declining population growth rate, attributed to lower winter survival rates over the last 70 years in the southern portions of the SBW range, could explain the decreased propensity of SBW to defoliate these areas. We show that severe defoliation resulting from the ongoing outbreak, especially areas that have not been affected by the previous outbreak, occurred under climate conditions that were much more favorable for population growth than during the previous outbreak.

Our findings indicate that contemporary alterations in forest landscapes are unlikely to have significantly influenced the recent changes in outbreak locations. Extensive timber harvest and land-use changes were the most important drivers of northeastern North American forests for several decades (Abrams, 1998; Danneyrolles et al., 2019). Over the last 50 years, these alterations gradually translated into a decrease in old-growth coniferous forests to the profit of younger, mixed or deciduous stands in the eastern boreal forest (Dupuis et al., 2020; Marchais et al., 2022). Forest management practices that alter forest composition and connectivity are known to influence spruce budworm outbreak dynamics, with managed forests enclosing a higher proportion of non-host species and younger age classes also showing reduced SBW outbreak intensity and frequency (Robert et al., 2018). These stands are less likely to include most vulnerable host species and as such, are making landscapes globally less vulnerable to SBW outbreaks (MacLean et al., 2001). But as these shifts were rather evenly distributed throughout the study area, they are less likely to have contributed to spatial modifications in outbreak severity at the supraregional scale. As such, our findings contrast with studies that showed that recent climate-induced or anthropogenic changes in hosts has altered insect pest outbreaks (Bouget & Duelli, 2004; Kausrud et al., 2012). That being said, landscape vulnerability to SBW is still currently highest in regions that have been severely affected by the ongoing outbreak (data not shown). The reduced vulnerability of landscapes further south compared to 50 years ago may also account for the northern location of the current outbreak where forest landscapes are most vulnerable.

These results demonstrate that recent climate change may have already surpassed the impacts of land use change on the dynamic of an important insect pest in North American forests. Historically, land use changes were the most important driver of biodiversity loss worldwide (Pereira et al., 2024), including for insects (Atmore & Buss, 2023). However, recent climate change puts additive and synergistic pressures on insects (Oliver et al., 2016) representing the most significant driver of the recent decline in insect abundance and shifts in community structure, development, dispersal patterns and phenology (Baranov et al., 2020; Engelhardt et al., 2022; Halsch et al., 2021). The great importance of temperature variations on insect ecophysiology is certainly no stranger to the fact that insects are generally more sensitive to climate change and show less inertia to these changes than other taxa (Vermaat et al., 2017). Indeed, our results contrast with other studies conducted in the same area that concluded that land-use changes had much more impacts on forest landscapes (Danneyrolles et al., 2019) and woodland caribou (Morineau et al., 2023) than climate change during the last century. High importance of recent climate change on a boreal insect pest species is also striking as high-latitude species usually show broader thermal tolerance than tropical species and hence more resilience to climate change (Bonebrake & Deutsch, 2012). Further anthropogenic climate forcing should exacerbate these alterations in insect life-history, confirming climate change as the most important driver in this regard in the following decades (Pereira et al., 2024).

Indeed, the northward shift in suitable climate conditions for SBW we estimated over the past 70 years serves as a prelude to projections under increasing anthropogenic climate forcing through the end of the 21st century (Boulanger et al., 2016). Previous studies (Boulanger et al., 2016; Candau & Fleming, 2011; Régnière et al., 2012) have shown that warmer temperatures associated with climate change would shift suitable conditions further northward throughout the SBW range. Other forest insect pest species in North America (e.g., Mountain pine beetle, Safranyik et al., 2010; Spongy Moth, Régnière et al., 2009) were also projected to have their climate envelope shift poleward or to higher elevations as a result of longer seasons for larval development or warmer winters. Although these shifts could lead to more severe insect pest outbreaks or result in impacts emerging in novel regions (Fält-Nardmann et al., 2018), climate-induced changes in the SBW climate envelope might in fact decrease the overall severity of future outbreaks due to a mismatch between suitable climate conditions and vulnerable host species (Boulanger et al., 2016). Yet, climate-induced changes in synchrony between host budburst and larval emergence in spring could increase the vulnerability of black spruce and hence impacts on forest landscapes in the upcoming decades (Pureswaran et al., 2015). Recent surveys do locally report high mortality in black spruce stands following severe defoliation in northern portions of the ongoing outbreak (Bognounou et al., 2017). Our assessment of recent climate-induced trends in suitable conditions for SBW highlights the need for close monitoring of current and future impacts of SBW in boreal forest landscapes.

Although it was not studied here, climate change and shift in SBW distribution and tree hosts attacked could also affect the third trophic level, i.e., natural enemies. Previous studies have shown that successful parasitism and survival of *Tranosema rostrale* (Brischke) (Hymenoptera: Ichneumonidae), an important parasitoid species attacking low density SBW populations, was reduced with increased temperatures (Seehausen et al., 2017, 2018). Similarly, recent modelling on three SBW parasitoid species have shown that these species should shift northward with climate change (Régnière et al., 2020, 2021a, 2021b). However, the trophic interactions are more complex than just climatic suitability and considering that most SBW parasitoids are multivoltine and need alternate hosts during the year, the impact of climate change on mortality inflicted by parasitoids is difficult to predict.

Already noticeable impacts of climate change on forest insect pests as demonstrated in this study reinforce the necessity of quickly adapting forest management strategies to shifting and novel climate-induced threats to forest ecosystems (Mina et al., 2022). Shifting climate conditions for forest pests could affect forest resilience (Pureswaran et al., 2015) and have tremendous impacts on the forest sector (Dukes et al., 2009). Modifications in pest dynamics and their impacts could cumulate with climate-induced change in other natural disturbances (e.g., wildfires) to put further pressure on forest ecosystems and the forest sector (Boulanger & Pascual Puigdevall, 2021). As we navigate these uncertain times, a proactive and integrated approach in forest management and ecological research will be crucial to mitigate the compounded effects of climate change and preserve the integrity and sustainability of forest ecosystems (Millar et al., 2007).

## Notes

### Competing Interest Statement

The authors have declared no competing interest.

